# Deep population structure linked to host vernalization requirement in the barley net blotch fungal pathogen

**DOI:** 10.1101/2023.12.20.572564

**Authors:** Julie Ramírez Martínez, Sonia Guillou, Stéphanie Le Prieur, Pauline Di Vittorio, Florelle Bonal, Demetris Taliadoros, Elise Guéret, Elisabeth Fournier, Eva H. Stukenbrock, Romain Valade, Pierre Gladieux

## Abstract

Invasive fungal pathogens pose a substantial threat to widely cultivated crop species, owing to their capacity to adapt to new hosts and new environmental conditions. Gaining insights into the demographic history of these pathogens and unraveling the mechanisms driving coevolutionary processes are crucial for the development of durably effective disease management programs. *Pyrenophora teres* is a significant fungal pathogen of barley, consisting of two lineages, Ptt and Ptm, with global distributions and demographic histories reflecting barley domestication and spread. However, the factors influencing the population structure of *P. teres* remain poorly understood, despite the varietal and environmental heterogeneity of barley agrosystems. Here, we report on the population genomic structure of *P. teres* in France and globally. We used genotyping-by-sequencing to show that Ptt and Ptm can coexist in the same area in France, with Ptt predominating. Furthermore, we showed that differences in the vernalization requirement of barley varieties were associated with population differentiation in France and at a global scale, with one population cluster found on spring barley and another population cluster found on winter barley. Our results demonstrate how cultivation conditions, possibly associated with genetic differences between host populations, can be associated with the maintenance of divergent invasive pathogen populations coexisting over large geographic areas. This study not only advances our understanding of the coevolutionary dynamics of the Pt-barley pathosystem but also prompts further research on the relative contributions of adaptation to the host versus adaptation to abiotic conditions in shaping Ptt populations.

**Impact statement:** Many invasive fungal pathogens have successfully followed major crop species throughout their intercontinental range, but continue to represent dynamic biotic threats. During their geographic expansion, invasive fungal populations were subjected to heterogeneous environmental conditions, or different populations of hosts, which could result in adaptation processes. Understanding this history of colonization can allow us to better prevent the emergence of infectious diseases of crops, and to better control them.

One such fungus, *Pyrenophora teres*, negatively impacts barley production globally by causing net blotch disease. In this study, we characterized the genetic makeup of *P. teres* in France and how it compares with what can be sampled in other regions of the world. We found that both the net and spot forms of *Pyrenophora teres* can be in the same area in France, but the spot form is more common. We also discovered that the net form populations associated with winter and spring barley are different, which was not known until now. This study opens up numerous experimental perspectives aimed at evaluating whether the two populations of net form are adapted to their hosts or to the conditions of cultivation of their hosts, with the goal of implementing measures that force the pathogen to maladaptation.

*Data summary:* GBS data are available under BioProject PRJEB66440. Single nucleotide polymorphism and reference genome assembly are available under doi: https://doi.org/10.5281/zenodo.10021844. Reads used for genome assembly are available under the BioProject PRJEB66476. The authors confirm all supporting data, code, and protocols have been provided within the article or through supplementary data files.

## Introduction

Numerous prominent fungal pathogens that affect major crops exhibit extensive geographic ranges [1–3]. These introduced fungal pathogens, which now thrive on physiologically- and phenologically-homogenous hosts planted over vast areas, have reached their current distribution following the colonization of a variety of environments, with demographic histories strongly constrained by the actions of their human vectors [4, 5]. As fungal pathogens established in new territories, they encountered new environmental conditions, new wild or domesticated host species, or new host varieties, potentially leading to local adaptation [6–9]. In ascomycetes, adaptation to new hosts (*i.e.*, specialization) is often associated with the differentiation of new lineages or species, which is expected for pathogens that use senescent host tissues as a substrate for reproduction, with no or little migration of gametes between infection and mating [10–12]. The relative weakness of prezygotic reproductive barriers in plant pathogenic ascomycetes [13] may also facilitate the emergence of local adaptation through hybridization or the introgression of adaptive variants [14–17]. Molecular population genetic analyses of whole-genome polymorphism data can reveal the demographic history of invasive fungal pathogens, provide insight into the mechanisms underlying coevolutionary processes [4, 18–22], and thereby help the implementation of durably efficient disease management programs [23–25]. Understanding how crop diseases spread, adapted, and co-evolved with their hosts requires a comprehensive picture of the evolutionary trajectory of fungal pathogens in conjunction with the (often complex) history of breeding, cultivation, and trade of their hosts [7, 26–29].

*Pyrenophora teres* is a major fungal pathogen of barley, present in all areas of barley cultivation [30–33]. *Pyrenophora teres* exists as two distinct lineages (called forms) that cause different symptoms. *Pyrenophora teres* form *teres* (Ptt) produces net-like lesions while *P. teres* f. *maculata* (Ptm) causes brown spots surrounded by chlorosis, thus referred to as “net form net blotch” and “spot form net blotch”, respectively. Ptt and Ptm cross easily in lab conditions [34], their artificial hybrids are fertile [35], and they can produce clonal hybrid lineages in the field [36]. However, population genomic studies revealed neither pervasive admixture nor rampant gene flow [37–40], which suggests that ecologically-based reproductive barriers contribute to the maintenance of Ptt and Ptm in syntopy on the same host [41]. Ptt and Ptm diverged well before the onset of plant domestication and agriculture, likely in different regions and/or on different hosts, with relatively recent secondary contact on barley [42, 43]. Although Ptt and Ptm both have global distributions, their relative abundance changes over time and space, with one form generally predominant in a given region [44–46]. The genetic structure of Ptm has been elucidated regionally, but its global structure remains unknown [37, 47–49]. Analyses of whole-genome sequencing data revealed that Ptt is structured into different populations, some tending to be geographically restricted to different regions, while others are distributed across multiple continents [29, 50]. Demographic modeling showed that the population structure of Ptt was shaped by the history of domestication and worldwide spread of barley. However, besides the contingencies linked to the history of the host and their impact on population demography, the nature of the host features responsible for the structure of Ptt remains poorly known despite the significant spatiotemporal heterogeneity of the barley agrosystem. Barley is an ancient crop that adapted to a broad spectrum of agricultural environments during a process of widespread range extension [51]. Domesticated barley is genetically structured into different populations, and the observed associations between agronomically-relevant phenotypes and population structure suggest that selection has played a role in the origin and/or maintenance of population differentiation [52, 53]. In particular, domesticated barley populations can vary in terms of cold requirement, with winter barley that requires lower temperatures for vernalization, and can therefore be sown in winter or late fall, and spring barley that does not require low temperatures and can be sown in spring. Domesticated barley populations can also vary in terms of grain characteristics, which is reflected in the morphology of ears, with six-row barley more often used as an animal or human feed, and two-row barley favored for malting and brewing. Some regions, such as France, have varied environmental characteristics, which allow the cultivation of different types of barley over large areas, providing ideal conditions to study the interplay between barley host diversity and the population structure of the pathogen. Here, we describe the population genetic structure of *P. teres* in France by using genotyping-by-sequencing (GBS) data. These data were used (1) to assess what are the forms of *P. teres* associated with barley crops in continental France and what is their distribution, (2) to investigate the contribution of various invasive populations from other continents to the genetic makeup of *P. teres* in France, and (3) to identify the host features associated with population differentiation.

## Methods

### Sampling, isolation, DNA extraction, and genotyping-by-sequencing

Random sampling of symptomatic barley leaves was performed from 2018 to 2021 in barley nurseries and experimental fields. A total of 16 localities in 12 administrative divisions (*i.e.*, *départements*) in France were included, and samples from several varieties and different barley types in terms of vernalization and row number were obtained. After sampling, the symptomatic tissue was either processed immediately for single-conidium isolation, or preserved at 4°C until processing. Monosporic cultures were obtained by three steps: (i) a colony was obtained by the inoculation of a PDA Petri dish with one conidium obtained from the symptomatic tissue, (ii) a quick DNA extraction was made using Chelex 100 by Bio-Rad, and the DNA was used to make a PCR using the primers and methodology developed by Leisova et al. [77] to screen for the presence of *Pyrenophora teres*, then, (iii) for the colonies in which the presence of the pathogen was confirmed, a single conidium was taken to produce new colonies that were used in the rest of the study. Alternatively, because some of the strains did not produce any more conidia after being obtained from the leaf, isolation was performed by cutting and transferring the tip of a hyphae.

DNA extraction was performed following the methodology proposed by Carlsen et al. [33] with some modifications. For each isolate obtained, five plugs were inoculated into a 250 mL Erlenmeyer containing 60 mL of modified Fries medium [33] , which was incubated at 27°C and 120 rpm for seven days. The content of the Erlenmeyer was blended and mixed with an additional 60mL of Fries medium and incubated for two more days under the same conditions. After incubation, the mycelium was rinsed with sterile distilled water and recovered using a piece of veil fabric as the filter (diameter of pores <0.3mm). The liquid was removed by pressing the mycelium contained in the fabric with the help of paper towels. The mycelium was frozen at -20°C and lyophilized during 24-36h, depending on the amount of humidity. The lyophilized tissue was placed in a 2mL microcentrifuge tube and ground using liquid nitrogen and a drill with plastic blue pellet pestles (Sigma-Aldrich). Just the sufficient amount to reach the line of 200µL of the tube was used for the extraction, following the protocol described by Carlsen et al. [33] but adding an extra RNAse treatment. The quality of the DNA was assessed using Nanodrop and electrophoresis on agarose gel, and the quantity was measured using Qubit. The DNA samples were stored at -20°C until used.

Genotyping-by-sequencing (GBS) libraries were constructed using 100ng of DNA for each isolate. DNA was digested with the ApeKI enzyme. After the ligation reaction, barcoded samples (5µL of each) were pooled and DNA was amplified. Purification was performed to remove adaptors and primer dimers using the Wizard Genomic DNA Kit (Promega) and quality was verified by measuring the quantity and length of the fragments. Libraries were sequenced using NovaSeq 6000 Illumina technology. Sequencing reads were demultiplexed using GBSX 1.3 [78], and trimmed for barcodes and adaptors using TRIMMOMATIC 0.39 [79]. Reads were inspected with FASTQC 0.11.9 and MULTIQC 1.11 [80, 81].

### Assignment of isolates to the two forms of *Pyrenophora teres*

For taxonomic affiliation, GBS reads were mapped to a reference genome of each of the possible forms of *P. teres* [54, 55] using BOWTIE 2.5 [82] with default parameters. SNP calling and filtering were performed using BCFTOOLS 1.16 [83]. After plotting for each individual, using histograms, the distribution of mapping quality (MQ) and sequencing depth, filtering cutoffs were set at MQ ≥40, and sequencing depth by individual ≥3. Sites with more than 50% missing data were removed using VCFTOOLS 0.1.16 [84]. The Variant Calling Files (VCFs; one file with Ptt as the reference, and one file with Ptm) were converted into FASTA pseudo-alignment files that were used to build neighbor-net phylogenetic networks using SPLITSTREE 4.17.2 [85]. Isolates were assigned to the reference genome to which they were the closest based on the p-distance (*i.e.*, Hamming distance).

Divergence between the two forms was quantified by computing net divergence as D_a_= d_XY_ - ( (π_Ptt_ +π_Ptm_ )/2 )), with d_XY_ representing nucleotide divergence and π representing nucleotide diversity. The mapping of short reads from Ptt on a Ptm reference, and vice versa, resulted in a significant underestimate of nucleotide diversity in both forms (not shown); it was therefore not possible to estimate the net divergence between the two forms using SNP calling data. We thus used public genomic data for six isolates of each form of *P. teres* (Supplementary table 1) to estimate D_a_ from the analysis of single-copy sequences identified with BUSCO 5.5.0 [86] after masking the genomes using REPEATMASKER v4 [87] with repeat families obtained with REPEATMODELER 2.0.3 [88]. The sequences of the 5551 single copy genes identified in all twelve genomes were individually aligned using MAFFT 7 [89], and d_XY_ and π were computed using the EGGLIB 3.3.0 [90] python package.

### Population subdivision in Ptt

To infer population subdivision in Ptt, we sequenced the whole genome of one of the isolates and used the assembly as the reference genome for calling SNPs in aligned GBS reads. The DNA of isolate FRA0042 was used to build a library with an insert size of 550bp, using the TruSeq Nano DNA Library Preparation Kit (Illumina). The library was sequenced on an Illumina NovaSeq 6000 machine. Sequencing reads were inspected and trimmed using the same approach as with the GBS data, and assembled using different K-mer sizes using ABYSS [91]. The chosen K-mer size was the one that maximized N50, L50, and assembly size (supplementary table 2). Quality of the resulting assembly was assessed using BUSCO after masking the repeats using REPEATMODELER and REPEATMASKER. GBS reads were mapped onto the reference genome using BOWTIE 2.5 [82] with default parameters, and filtered using BCFTOOLS 1.16 [83]. Filtering cutoffs were set at MQ ≥50, and sequencing depth by individual ≥3, after examination of the distribution of these statistics for each individual. The single-nucleotide polymorphisms (SNPs) used in the analysis of population subdivision were obtained by filtering out indels, monomorphic sites, and sites with more than 50% missing genotypes.

Population subdivision was inferred in R 4.2.1 using approaches that make no assumption of linkage or Hardy-Weinberg equilibrium: the clustering algorithm SNMF (sparse non-negative matrix factorization) implemented in the package LEA [92], and DAPC (Discriminant Analysis of Principal Components) implemented in the package POPPR [93]. The host features and other factors associated with population differentiation were investigated using Principal Component Analysis (PCA) and Analysis of Molecular Variance (AMOVA) using the packages POPPR and ADEGENET in R. The factors considered were the barley type, year of sampling, barley variety, site of origin, and the clusters identified with clustering algorithms.

### Summary statistics of nucleotide variation

The dataset used in the computation of summary statistics was obtained by filtering out indels from the VCF produced after filtration using BCFTOOLS (i.e., sites with missing data and monomorphic sites were kept in the dataset). Nucleotide diversity π, and nucleotide divergence d_XY_ were estimated in 100Kbp windows using the program PIXY 1.27b [94] , which provides unbiased estimates of diversity and divergence in the presence of missing data. Linkage disequilibrium decay (LDD) was assessed using the package POPLDDECAY [95], by generating a pseudo-diploidized VCF.

### Global population structure of Ptt

The evolutionary relationships between the French isolates of Ptt and populations from other countries were determined by combining our GBS data with previously released whole-genome sequencing data [29] , and with newly generated GBS for 15 isolates from Canada (n=6) [37] and USA (n=9). The fifteen isolates were grown, processed for DNA extraction, and genotyped with the same methodology as described above for French isolates. Sequencing reads were mapped onto the Ptt reference genome [55] and the SNP calling was performed as already described above for GBS data. The VCF was filtered by removing indels, and keeping only biallelic variants with less than 5% missing genotypes. The final file contained samples from the USA (n=29), Canada (n=6), Morocco (n=27), Iran (n=9), Azerbaijan (n=21), Denmark (n=5), and more samples from France (n=204 with GBS data [this study] and n=21 with whole-genome data [29]) (Supplementary table 3). Population subdivision was analyzed making a PCA using the package POPPR in R.

## Results and discussion

### Ptt outpaces Ptm in frequency as a barley pathogen in France

A total of 207 isolates of *Pyrenophora teres* were obtained from four collection campaigns in 2018 (n=39), 2019 (n=11), 2020 (n=94), and 2021 (n=63). Isolates represented 16 localities distributed in 12 administrative divisions of France (i.e., *départements*), and more than 15 barley varieties of two-row spring barley (n=26) and winter barley (n=122). All but two of the winter barley varieties were of the six-row type. Information about the type of barley (winter *vs.* spring) was available for all 207 isolates and the identity of the variety of origin was available for 204 isolates. Sampling information is provided in Supplementary table 4.

The GBS reads covered 30 and 32%, respectively, of the reference genomes of Ptt and Ptm [54, 55] (Supplementary table 5). SNP calling and filtering identified 1.1e5 SNPs for the dataset with Ptt as the reference genome, and 8.4e4 SNPs for the dataset with Ptm as the reference genome. Neighbor-net networks built from these datasets showed that most of the isolates (n=204) were closely related to the reference genome of Ptt (Figure 1A), with a p-distance from the Ptt reference genome of 0.08 differences/polymorphic site on average. Only three isolates were closely related to the reference genome of Ptm (Figure 1B), with a p-distance from the Ptm reference genome of 0.2 differences/polymorphic site on average (Supplementary tables 6 and 7). Net divergence (D_a_) between the two forms, as estimated from alignments of single-copy orthologs identified in publicly available genomes, was 0.005/bp (s.d. 0.016) (Supplementary table 8), which can be used to estimate that the two forms diverged 1.1e5 years ago, assuming a substitution rate of 2.2e-8 per base pair [56, 57]. The estimated level of net divergence between the two forms of *P. teres* places them in the gray zone of speciation (0.5-2% net synonymous divergence), in which taxonomy is often controversial, and hybridization possible [58]. However, the absence of reticulations along the branches connecting Ptt and Ptm indicated the absence of hybrids in our dataset (Figure 1A-B). This suggests that intrinsic postzygotic barriers (*i.e.*, incompatibilities [41]), as well as possible extrinsic prezygotic barriers that remain to be identified (*e.g.*, adaptation to the host [6, 59]), may contribute efficiently to the maintenance of the two forms on the same crop species without pervasive hybridization.

**Figure 1.**
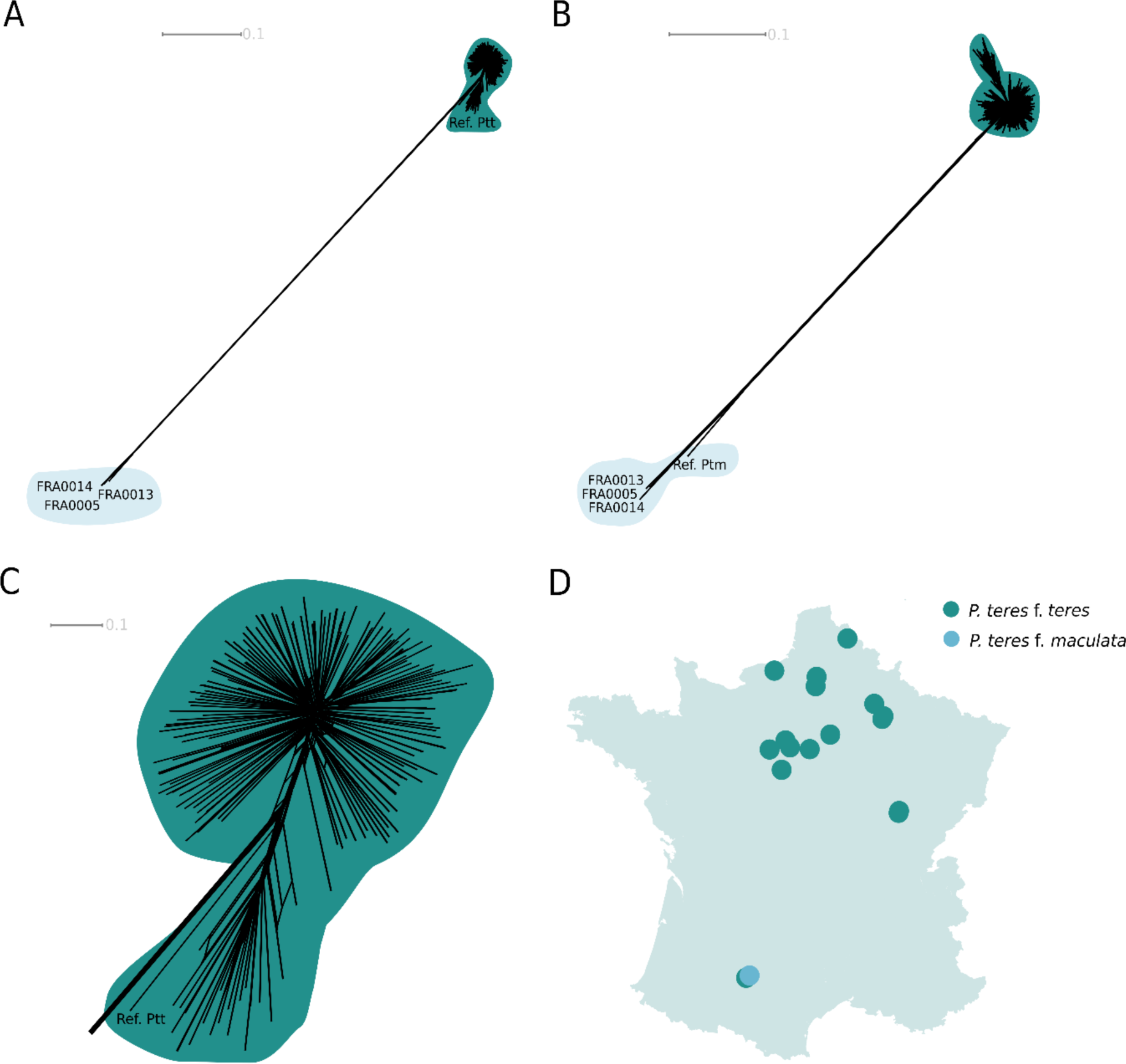
Identification of the two different forms of *Pyrenophora teres* present among isolates from France and their geographic distribution inferred by the analysis of neighbor-net networks generated from genotyping-by-sequencing (GBS) data. Two main groups are highlighted, the green group showing isolates identified as Ptt, and the light blue group showing isolates identified as Ptm. Isolate names are not shown for the Ptt group. **A**. Network constructed from the dataset obtained by mapping GBS reads onto the reference genome of *Pyrenophora teres* f. *teres* (Ptt), which is referred to as “Ref. Ptt”. **B**. Network constructed from the dataset obtained by mapping GBS reads onto the reference genome of *Pyrenophora teres* f. *maculata*, which is referred to as “Ref Ptm”. **C**. Closer view of the group formed by isolates that were the most closely related to the Ptt reference genome. **D.** Map of geographic distribution of Ptt and Ptm in France. Dots correspond to sampling sites, and their color to the different taxa found at each site. The two dots at the bottom left of the map correspond to the same geographical site; one of the two dots was moved slightly to avoid complete overlap. The p-distance scale bar (in number of differences per polymorphic site) is displayed at the top of panels A, B, and C.

We show that Ptt is largely dominant over Ptm in France, with only 1.4% of isolates assigned to Ptm, which suggests there was a swap during the past three decades since the most prevalent form of *P. tere*s in France 30 years ago was Ptm [60]. The study of this pathogen in other regions showed that the prevalence of the forms can rapidly change depending on control strategies like the presence of resistant plant material [61, 62]. Our findings suggest that the prevalence of the two forms in France could change again in the future, highlighting the need for regular monitoring of *P. teres* populations.

Ptt was present in all of the sampled areas in France, whereas Ptm was only found in the southwest of the country, where it coexisted with Ptt in the same field. This pattern is similar to what has been reported in other regions of Eastern and Northern Europe, North America, Italy, the Maghreb, and Egypt, where the two forms are present, but Ptt clearly predominates [37, 46, 59, 62–72]. Our results, however, contrast with what was observed in East Africa, South Africa, and Southeastern Australia, where Ptm is more prevalent [44, 61, 73]. The differences in prevalence could be explained by environmental conditions, or by the heterogeneity in the distribution of varieties with differences in levels of resistance to the two net blotch pathogens [59]. Based on our sampling, we cannot favor one hypothesis over the other due to the low prevalence of Ptm. Our findings do indicate, however, that it is possible for both forms to coexist on different hosts in sympatry, which opens up opportunities for future experiments to test the hypothesis that the two forms of net blotch pathogens are adapted to different populations of barley.

### *Pyrenophora teres* f. *teres* is subdivided into two main clusters in France

The population structure of the 204 Ptt isolates was investigated by mapping GBS reads onto the assembled genome of the isolate FRA0042. The genome assembly of FRA0042 consisted of 6.8e4 contigs larger than 1 Mb, and was characterized by an N50 of 328Kb, an L50 of 30, and a BUSCO score of 94.4%. On average across isolates, 43% of the 32 Mb reference genome was covered by GBS reads (Supplementary table 5). The final filtered dataset included 6.3e4 SNPs.

Analyses with the SNMF algorithm and Discriminant Analysis of Principal Components (DAPC) identified two well-differentiated clusters within Ptt. In analyses with SNMF, cross-entropy decreased monotonically with increasing K values (Supplementary figure 1 and Supplementary table 9). However, only the model with K=2 clusters showed two well-separated groups (Figure 2A). Increasing K did not identify clear new clusters, and this simply resulted in a breakup of ancestry proportions between different clusters. In the DAPC, the minimum value of the Bayesian information criterion (BIC) was observed at K=3 (Supplementary figure 1), but only K=2 presented a clear pattern of subdivision (Figure 2). Clusters identified in models with K≥3 exhibited significant overlap along the second discriminant function (Figure 2E; Supplementary Figure 1). Analysis of molecular variance revealed that differentiation was relatively high between clusters (Supplementary table 10), with 45% of the total variation being distributed between clusters. Net divergence, which scales linearly with time under a strict isolation model [57], was D_a_=0.0014 between the two clusters, which represents 20% of the estimated net divergence between Ptt and Ptm (Supplementary table 11). Despite the recent divergence between clusters, admixture appeared limited, with only 0.5% of isolates presenting ancestry proportions between 0.2 and 0.8 in analyses with SNMF (Supplementary table 4).

**Figure 2.**
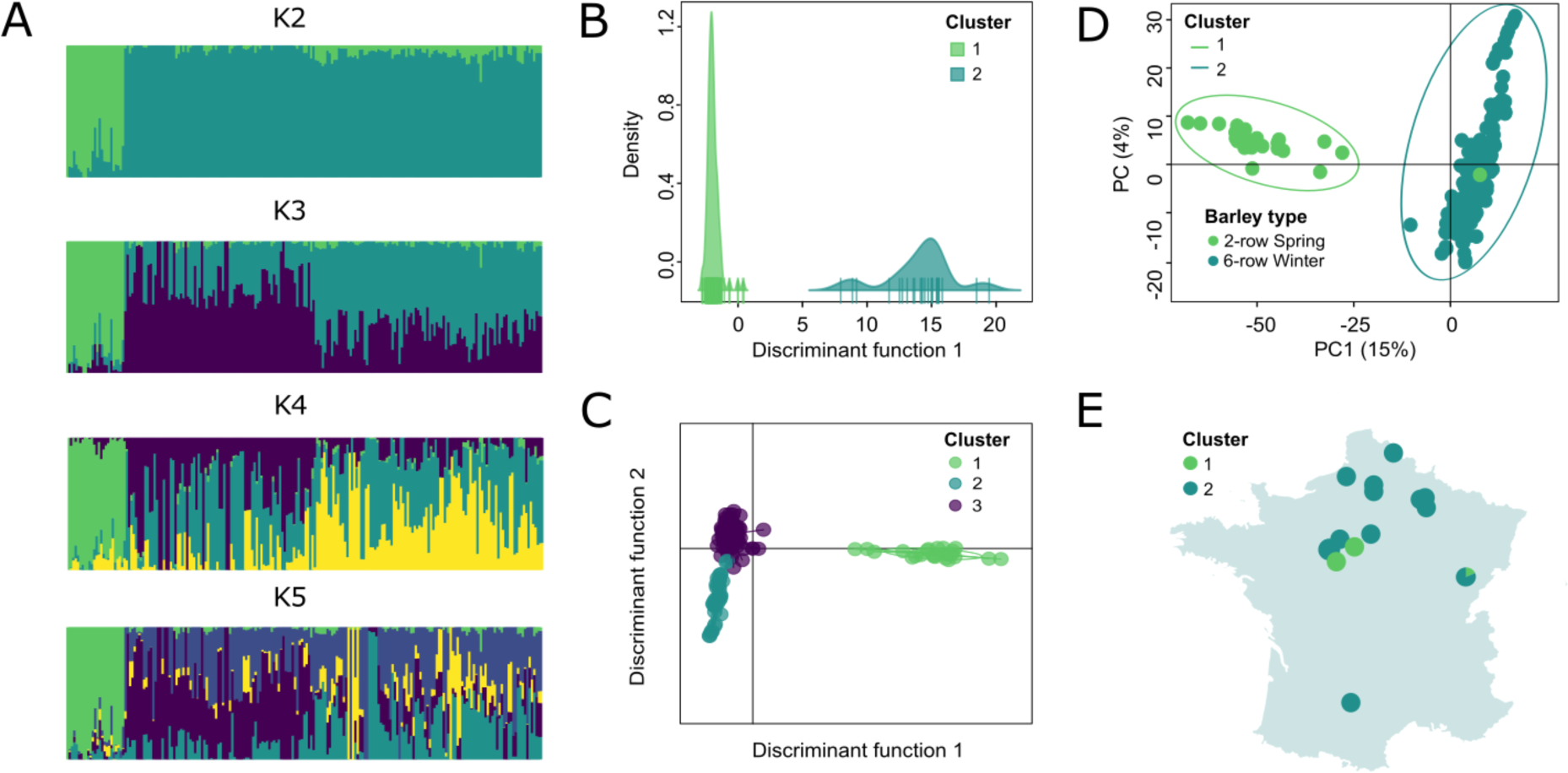
Population subdivision in *Pyrenophora teres* f. *teres* from France, estimated from genotyping-by-sequencing data for 204 isolates from 16 sampling sites. **A**. Ancestry proportions in K clusters estimated with sNMF, with each isolated represented by a segment subdivided in K intervals. **B**. Discriminant Analysis of Principal Components (DAPC) with two clusters, based on 20 principal components. **C**. DAPC with three clusters, based on 25 principal components. **D.** Principal Component Analysis of genotype data, with each isolate represented by a dot colored based on its type of barley of origin, and ellipses showing clusters of isolates identified using DAPC and sNMF analysis. **E.** Geographical distribution of the two main *Pyrenophora teres* f. *teres* clusters in France, with pie charts representing the proportion of isolates from each cluster at each sampling site.

The two main clusters identified in Ptt displayed similar genetic structures. Summary statistics were calculated based on 2.5e6 sites passing quality filters. The two clusters displayed close estimates of nucleotide diversity (cluster 1: π=0.00434; cluster 2: π=0.00433), despite differences in Tajima’s D suggesting contrasted demographic histories (cluster 1: D=1.47; cluster 2: D=-0.72; Supplementary table 11). Patterns of linkage disequilibrium decay were consistent with a history of recombination, and thus sexual reproduction, both at the scale of Ptt and clusters within Ptt, with linkage disequilibrium reaching half of its maximum value within less than 5e3 bp (Supplementary figure 2). Analyses of molecular variances revealed limited sub-structure within clusters, with more than 90% of variation distributed within locations or varieties (Supplementary table 12), which is consistent with the lack of clear subdivision observed in models with K>=2 clusters with the SNMF algorithm and the DAPC. Together, these analyses indicate a history of recent and widespread gene flow within clusters, and therefore the lack of strong barriers to dispersal and the absence of lineages strongly differentiated by the effect of local adaptation to host varieties.

The two major clusters identified within French Ptt were associated with different types of barley. We relied on PCAs to visualize the relationship between population structure and the isolates’ metadata, *i.e.*, barley variety, barley type, year of sampling, and site of origin. We found that the type of barley was the factor most clearly associated with population subdivision, with the first cluster being formed almost entirely by isolates from 2-row spring barley, except isolate FRA0049, and the second cluster grouping isolates from 6-row winter barley (Figure 2D; Supplementary figure 3, Supplementary tables 4 and 9). The two clusters were thus distributed according to the location of the types of barley sampled as shown in figure 2E. In our sampling, only one type of barley was generally sampled per location, and consequently, only one cluster was detected. Interestingly, there was only one area (*Côte d’Or*, in Burgundy) in which both clusters were present and no significant difference in the membership proportions estimated by SNMF between the isolates from this area and other regions was found, neither when assigned to cluster 1 nor to cluster 2 (Wilkoxon’s test p-value = 0.562 and 0.067 respectively, see Supplementary figure 4). This indicates that the two populations can coexist without any detectable increase in admixture levels, and adaptation to the host may play a role in maintaining these populations. However, grain type and vernalization requirement were associated in the barley varieties we sampled (i.e., all 2-row varieties were spring barleys, and all 6-row varieties were winter barleys, the only two samples from 2-row winter barley obtained were classified as Ptm), and we, therefore, cannot predict which of these two traits would be more likely to be associated with the differentiation between the two clusters in a more diverse barley panel.

### The global structure of Ptt is associated with the vernalization requirements of barley varieties

To determine where the French isolates branch into the global structure of Ptt, but also to further evaluate the association between barley type and population structure, we combined our GBS dataset with publicly available whole-genome data [29]. The combined dataset, including 133,609 SNPs without missing data, was submitted to PCA. Principal components (PCs) 1, 2, and 3, represented 18, 8, and 5% of total variation, respectively. While PC2 did not reveal any evident association between differentiation and metadata (host or region of origin; Supplementary figure 5), PC1 and PC3 subdivided the dataset into several clusters that mirrored relatively well geographical origins (Figure 3A), consistent with previous findings [29]. PC1 identified three clusters: a first cluster mostly comprised of isolates collected in Azerbaijan and isolates collected in France from 6-row winter varieties, a second cluster mostly comprised of isolates from Iran, and a third cluster grouping the remaining isolates. PC3 split the third cluster identified along PC1 into one cluster mostly comprised of isolates from North America, one cluster mostly comprised of isolates from North Africa, and one last cluster mostly comprised of isolates collected from 2-row spring varieties in France. Our results thus demonstrate that what has previously been called the “Caucasus cluster” [29] is also present at a relatively high frequency in Europe.

**Figure 3.**
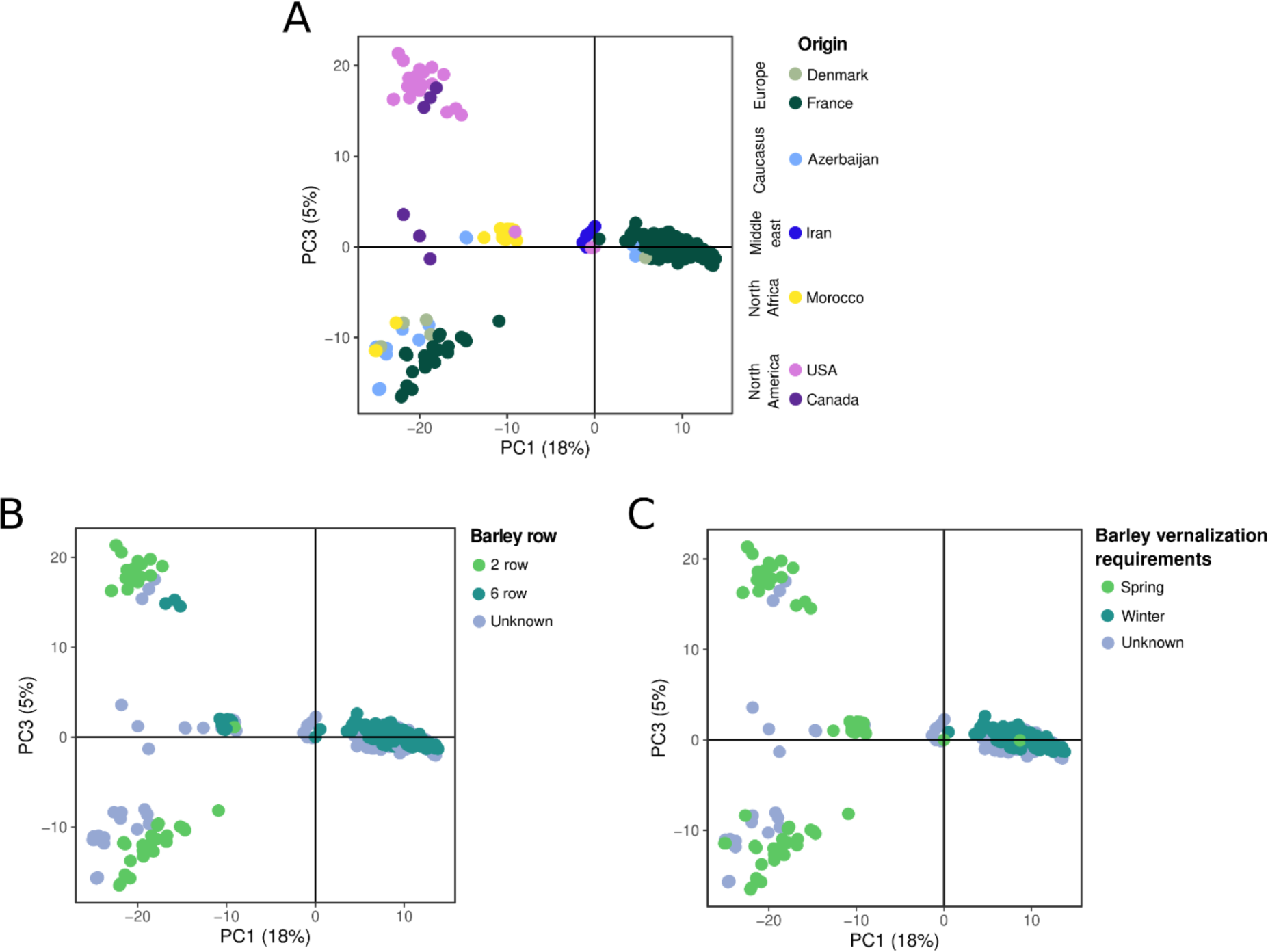
Association between population structure in *Pyrenophora teres* f. *teres* isolates collected globally, and the vernalization requirement of their barley host, as visualized using principal components analysis of single-nucleotide polymorphisms. Each dot represents an isolate. **A**. Isolates colored by country of origin. **B**. Isolates colored according to the number of rows of the barley variety from which they were obtained. **C.** Isolates colored according to the vernalization requirement of the barley variety from which they were obtained.

Although some geographical component to population structure was observed, another factor associated with differentiation within global populations of Ptt was the type of barley, and more precisely, the vernalization requirement. We were able to collect metadata for 46% of isolates from the entire genome dataset, which, when combined with the GBS isolates with known barley type, formed a new dataset where the type of grain and vernalization requirement were no longer strictly associated. This allowed us to independently assess the association of population differentiation with each factor using PCA. Although neither of the two principal components, PC1 and PC3 could distinguish between isolates from 2-row or 6-row barley (Figure 3B), PC1 was able to separate isolates from winter and spring barley in an almost mutually exclusive manner (Figure 3C). Our analysis does not contradict previous conclusions [29], but rather provides additional insights. Our results suggest that the Caucasian cluster, from which the spring barley population of North Africa originated, and from which the spring barley populations of North America and Europe diverged relatively long ago, is actually a population associated with winter barley and not specific to the Caucasus. This leads us to propose a hypothesis that adaptation to spring barley, or its growing conditions, would have played a crucial role in the emergence of Ptt.

## Concluding remarks

A growing body of evidence suggests that adaptation plays a major role in the successful emergence and dissemination of pathogens across heterogeneous environments in time and space [4, 7, 8, 12, 16, 74, 75]. In our study, we used a molecular ecology approach to show that there are genetic differences between populations of Ptt that are associated with winter and spring barley in France, as well as on a broader geographical scale. The existence of such a pattern suggests that the populations of the pathogen are differentially adapted to the two types of barley, and/or to the conditions in which they are cultivated. This work opens up numerous research perspectives to determine what are the barriers, particularly prezygotic [10, 76], that contribute to the maintenance of the two populations, and in particular, what is the relative contribution of adaptation to the host or adaptation to abiotic conditions. Our results also highlight the importance of testing new resistant barley materials with both spring and winter barley isolates of Ptt, especially in areas where both types of barley are used.

## Abbreviations

Pt: Pyrenophora teres
Ptt: Pyrenophora teres form teres
Ptm: Pyrenophora teres form maculata
GBS: genotyping-by-sequencing
DAPC: Discriminant Analysis of Principal Components
PCA: Principal Component Analysis
AMOVA: Analysis of Molecular Variance

## Acknowledgements

To Timothy Friesen for kindly sharing methods for DNA extraction for *P. teres,* and answering our questions about the biology of *P. teres*. We are grateful to Shaun Clare and Robert Brueggeman from Washington State University, and to Stephen Strelkov and Rosa-Ileana Strelkov from the Plant Molecular Pathology lab of the University of Alberta for sharing *P. teres* strains from the US and Canada. We thank Pierre Mournet at UMR AGAP in Montpellier for his help with GBS library construction, and Florent Remuson (ANSES) for isolating and providing us with isolates of *P. teres*.

