## Supplementary Figure 3 for "Deep population structure linked to host vernalization requirement in the barley net blotch fungal pathogen"

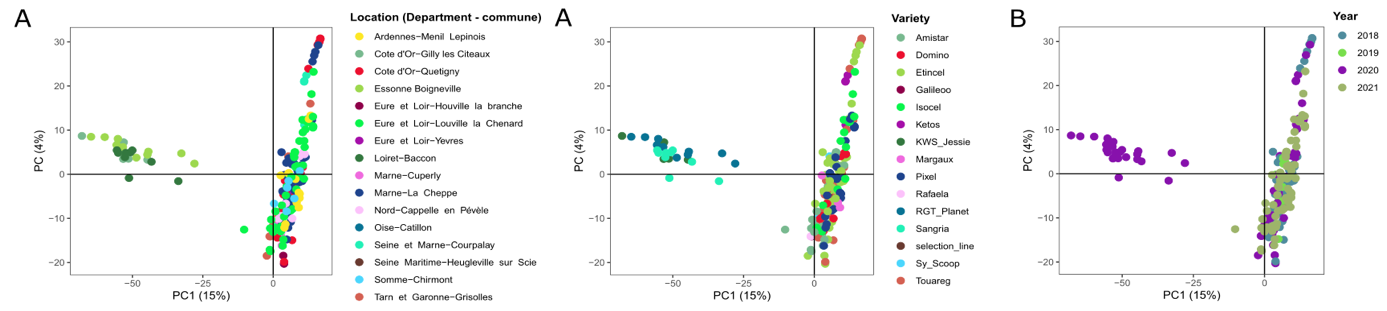

Supplementary Figure 3. Principal Component Analysis of genotype data of Pyrenophora teres f. teres associated to net blotch of barley in France, with each isolate represented by a dot. A. Isolates colored by location of sampling. B. Isolates colored according to barley variety from which they were obtained. C. Isolates colored according to the year of sampling.
