## Supplementary Figure 5 for "Deep population structure linked to host vernalization requirement in the barley net blotch fungal pathogen"

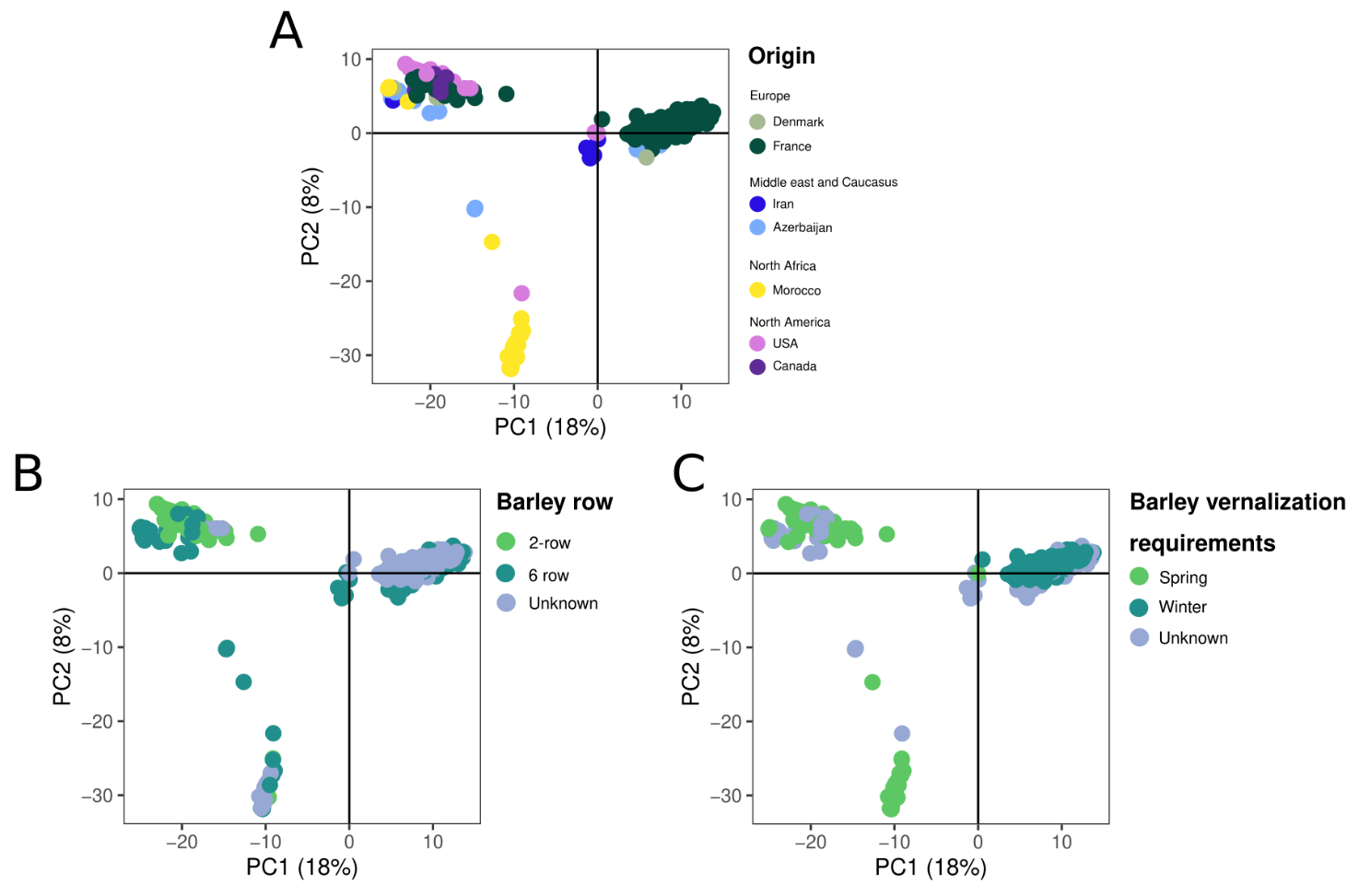

Supplementary Figure 5. Association between population structure in Pyrenophora teres f. teres isolates collected globally, and the vernalization requirement of their barley host, as visualized using principal components analysis of single-nucleotide polymorphisms showing PC1 and PC2. Each dot represents an isolate. A. Isolates colored by country of origin. B. Isolates colored according to the number of rows of the barley variety from which they were obtained. C. Isolates colored according to the vernalization requirement of the barley variety from which they were obtained.
