## Supplementary Figure 4 for "Deep population structure linked to host vernalization requirement in the barley net blotch fungal pathogen"

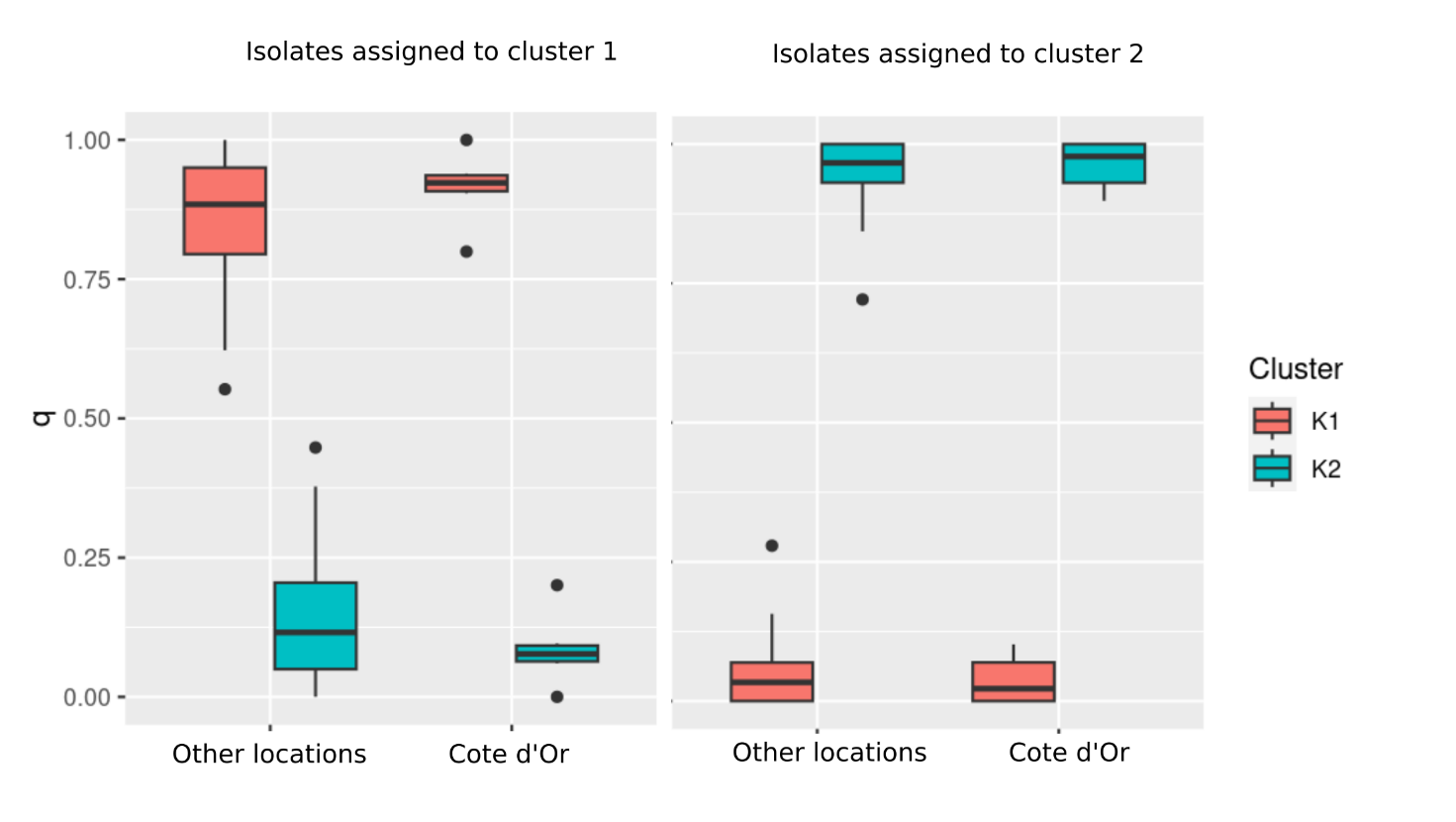

Supplementary Figure 4. Comparison of ancestry proportions (q) derived from sNMF analysis with K=2 for GBS data isolates of Pyrenophora teres f. teres in the Côte d’Or region of Burgundy versus other surveyed areas in France.
