## Supplementary Figure 2 for "Deep population structure linked to host vernalization requirement in the barley net blotch fungal pathogen"

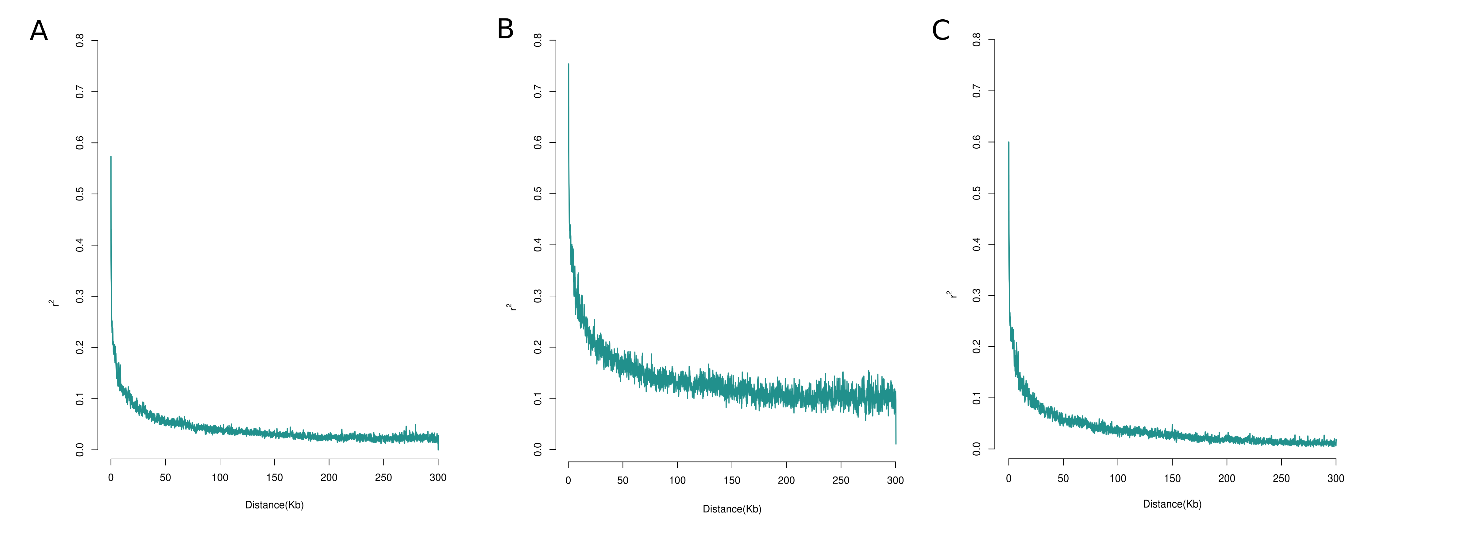

Supplementary Figure 2. Linkage disequilibrium decay shown as the squared correlations of allele frequencies (r2 ) plotted against distance between SNPs obtained from BGS data from Pyrenophora teres f. teres isolates from barley in France. A. Including all the isolates obtained (n=204) B – C. Including only the samples assigned to the different clusters found inside the isolates of Ptt. B. Including isolates assigned to cluster 1 (n=25). C. Including isolates assigned to cluster 2 (n=179).
