## Supplementary Figure 1 for "Deep population structure linked to host vernalization requirement in the barley net blotch fungal pathogen"

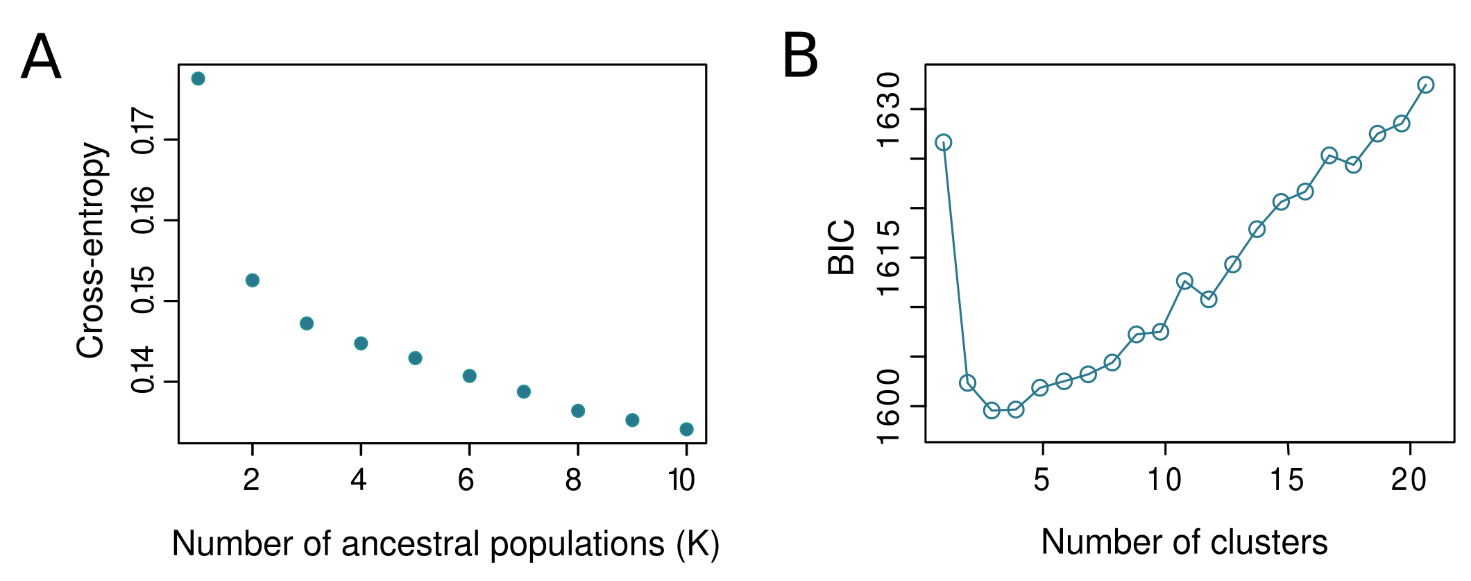

Supplementary Figure 1. Assessment of the number of clusters in the population of Pyrenophora teres f. teres isolates from barley in France. A. using cross- entropy values obtained by sNMF (non-negative matrix factorization) analysis. B. using Bayesian Information Criterion (BIC) of different clusters obtained by clustering analysis implemented in R.
